# Body-wrapping anterior flagella drive ultrafast swimming in bacterial zoospores

**DOI:** 10.64898/2026.03.29.715058

**Authors:** Naoki A. Uemura, Tsubasa Ishida, Yoshiyuki Sowa, Moon-Sun Jang, Takeaki Tezuka, Daisuke Nakane, Yasuo Ohnishi

## Abstract

Most bacteria swim at ∼10 body lengths per second (bl s^−1^), yet some microorganisms move far faster, and the physical design principles enabling such extreme motility remain poorly understood. Here we uncover an ultrafast swimming strategy in *Actinoplanes missouriensis* zoospores, which reach up to 560 μm s^−1^ (∼500 bl s^−1^), the fastest relative swimmer reported for any microorganism. High-speed imaging shows that propulsion is driven by an anterior, body-wrapping bundle of short flagella that rotate synchronously as a right-handed helix at ∼150 Hz. This architecture generates high thrust without extreme motor speeds and contrasts with the canonical rear-bundle paradigm of bacterial swimming. Microfluidic assays further demonstrate that this propulsion mode enhances dispersal across the flow interface, providing a mechanism for rapid colonization during the transient motile phase of the life cycle. Furthermore, chemotaxis-dependent reorientation is observed, suggesting that zoospore swimming can be directionally regulated. These results identify a new locomotion principle in which supramolecular organization of multiple flagella, rather than motor speed alone, sets the upper limits of bacterial swimming and offers inspiration for the design of high-performance microswimmers.

## Introduction

Who holds the title of the fastest swimmer on Earth? Across the tree of life, larger body size generally correlates with faster locomotion^1^. When swimming speed is normalized by body length^2^, most animals achieve speeds on the order of 10 body lengths per second (bl s^−1^). Some microorganisms, however, attain far greater values^3^ reaching up to 100 bl s^−1^, implying that distinct physical regimes and propulsion architectures can push the limits of locomotion.

Microorganisms exhibit a remarkable diversity of propulsion strategies adapted to low-Reynolds-number environments. Many bacteria and archaea, micron-sized organisms, thrust by rotating helical filaments, known as flagella or archaella^4–6^; however, the number of filaments and their spatial arrangement and higher-order organization vary widely and strongly influence performance. A representative bacterium, *Escherichia coli*, swims by rotating several flagellar filaments^7^ that bundle at the rear and drive the cell at ∼30 μm s^−1^, equivalent to ∼10 bl s^−1^. Other organisms achieve much higher speeds: magnetotactic coccoid bacteria swim at ∼300 μm s^−1^ (∼200 bl s^−1^) using tightly bundled flagellar filaments within a sheath^8,9^; hyperthermophilic archaea can reach ∼500 μm s^−1^ (∼400 bl s^−1^) at their optimal growth temperature^10^; and the bacterium *Candidatus* Ovobacter propellens utilizes a front bundle of hundreds of filaments to swim up to ∼1,000 μm s^−1^ (∼200 bl s^−1^)^11^. Given that the molecular machinery of bacterial flagella is largely conserved, these observations suggest that supramolecular organization sets the upper limits of swimming performance. What physical design principles enable such extreme speeds?

Actinomycetes are mainly soil-inhabiting bacteria, most of which have lost flagellar motility in favor of mycelial growth^12^. However, a subset retains flagella and produces motile spores or zoospores during a specific life stage^13,14^. Members of the genus *Actinoplanes* form terminal sporangia that release hundreds of spherical zoospores upon contact with water, allowing dispersal toward favorable environments before germination (Fig. 1a)^15^. The zoospore of *Actinoplanes missouriensis* possesses genetically characterized bacterial flagella^12,16^ and exhibits only a brief motile phase, swimming for ∼1 h before attaching to surfaces and initiating outgrowth, often aided by type IV pilus-mediated adhesion^17–19^.

**Figure 1:**
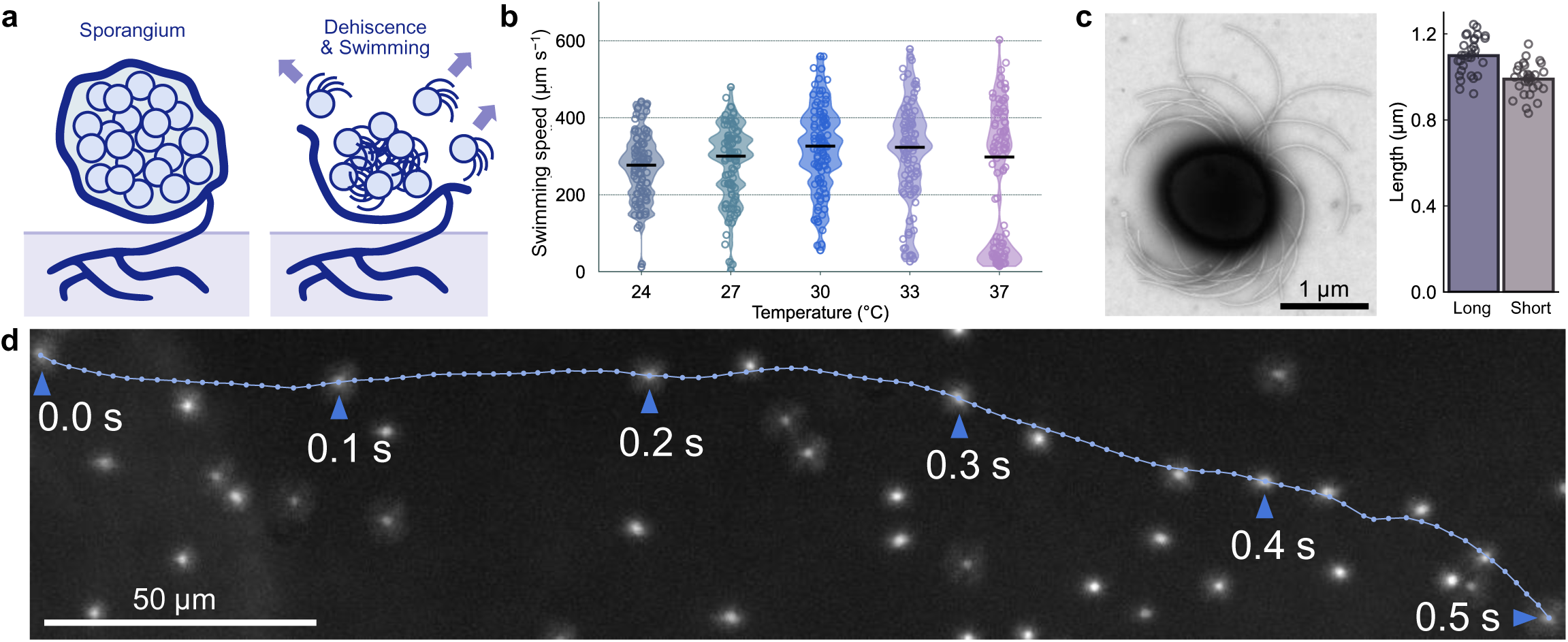
Ultra-fast swimming in zoospores of *A. missouriensis*. **a** Schematic illustration of sporangium dehiscence and release of swimming zoospores. **b** Swimming speed measured at different temperatures. Black bars indicate the median values, and circles represent biological replicates (*N* = 100 cells). **c** Size of zoospore. TEM image of zoospore (*left*) and distribution of cell diameters (*right*, *N* = 30 cells). **d** Swimming trajectory of a zoospore over 0.5 s at 30°C. Positions were recorded at 5-ms intervals. The trajectory is overlaid on a maximum-intensity projection of dark-field images acquired at 0.1-s intervals under dark-field microscopy. See also Movie S2.

Here we show that *A*. *missouriensis* zoospores achieve swimming speeds of up to ∼560 μm s^−1^ (∼500 bl s^−1^), representing the fastest relative swimming speed reported for any microorganism. High-speed imaging reveals that dozens of short flagellar filaments cluster at the anterior pole, wrap around the cell body, and rotate synchronously as a helical bundle at ∼150 Hz. We propose that this anterior body-wrapping architecture enables highly efficient thrust generation and represents a previously unrecognized design principle for ultrafast microscale locomotion.

## Results

### Ultra-fast swimming at optimal growth temperature

Although *A. missouriensis* zoospores are known to be motile, their swimming dynamics have not previously been quantified with high temporal resolution^14,15^. Using dark-field microscopy, we tracked individual zoospores immediately after release from sporangia and observed that they began swimming within seconds (Movie S1), indicating that their flagella are pre-assembled and fully functional at the moment of release. By optimizing sporangium dehiscence conditions and observation timing, we measured mean swimming speeds of 269 ± 88 μm s^−1^ at 24°C (Fig. 1b), exceeding previous reports^15^ of 135 ± 25 μm s^−1^. The trajectories were slightly helical, likely due to torque balance between the rotation of the flagella and the counter-rotation of the cell body. Swimming speed depended strongly on temperature, reaching 320 ± 112 μm s^−1^ at 30°C, which is the optimal temperature for mycelial growth of this bacterium (Fig. 1b and Movie S2), but declined at 37°C. Considering that the body size of zoospore is 1.1 ± 0.1 μm for the longer axis and 1.0 ± 0.1 μm for the shorter axis (Fig. 1c), these values correspond to ∼300 bl s^−1^, with some individuals reaching up to 560 μm s^−1^ (∼500 bl s^−1^) (Fig. 1d). These measurements establish the *A. missuriensis* zoospore as the fastest swimmer reported to date when normalized by body size.

### Flagellar-forward swimming drives ultrafast motion

To visualize flagellar filaments, we introduced a cysteine-substituted flagellin gene (*fliC*^S260C^) into a *fliC* null mutant (Δ*fliC*) strain using a chromosome-integrating vector, pTYM19-Apra^20,21^, enabling site-specific labelling with a maleimide-conjugated fluorophore^22^. In a previous study, the non-flagellated and non-motile phenotype of the Δ*fliC* strain was restored to the wild-type (WT) phenotype by the introduction of *fliC*^16^. The labelled mutant zoospores swam at speeds comparable to those of WT zoospores, demonstrating that this mutation did not affect swimming behavior (Fig. S1). Fluorescent labelling of flagellar filaments revealed that motile zoospores consistently position their flagella at the anterior pole, corresponding to the direction of movement (Fig. 2a *left* and Movie S3). During swimming, fluorescence signals remained confined to the anterior hemisphere, even in abrupt reorientation events (Fig. 2a *right*). After each turning event, cells resumed swimming with the same forward-facing configuration, demonstrating that the flagellar array remains structurally polarized toward the front. This arrangement contrasts with the posterior flagellar bundle observed in *E. coli*^7^.

**Figure 2:**
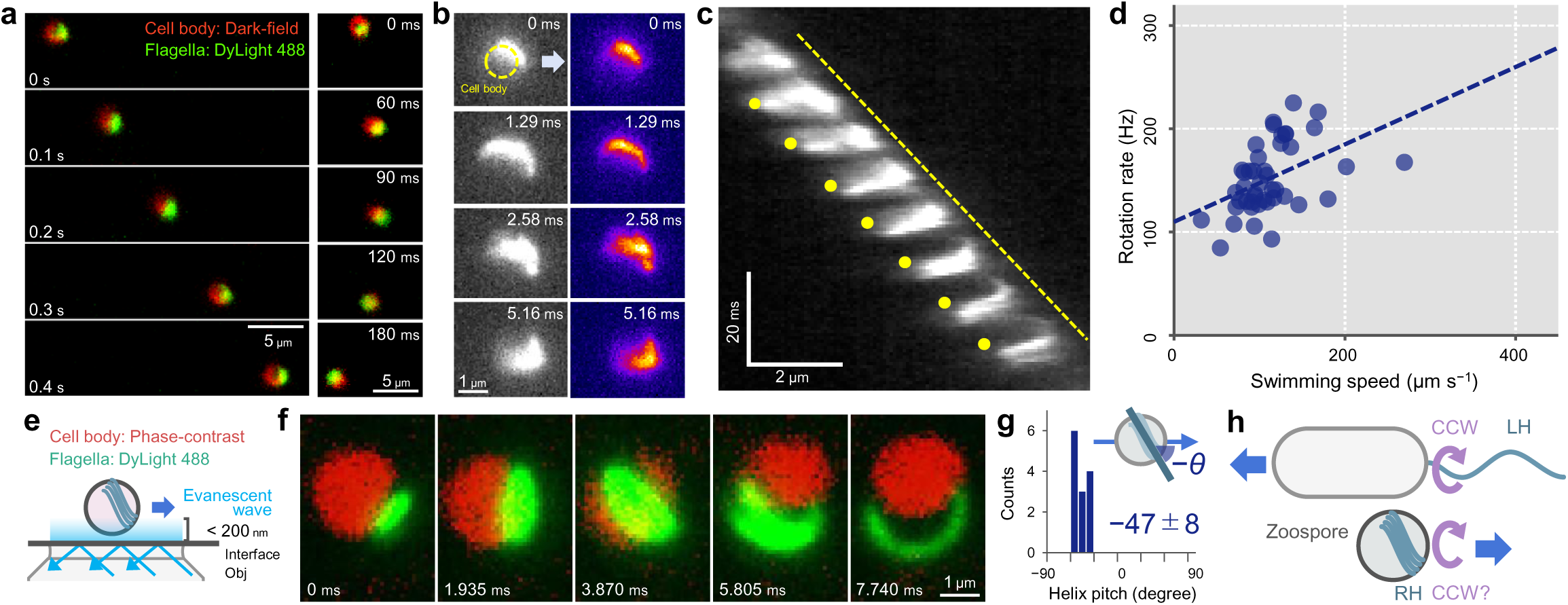
Anterior flagella propulsion in zoospore. **a** Sequential imaging of flagellar filaments during swimming. Flagellar filaments were fluorescently labeled and imaged at 10-ms intervals at 24°C. Swimming with anterior flagella (*left*) and directional reorientation (*right*). See also Movie S3. **b** High-speed images showing the rotation of the flagellar filament bundle at 24°C. Yellow circle indicates the cell body. See also Movie S4. **c** Kymograph of flagellar signals along the swimming axis. A diagonal dashed line represents cell displacement over time. The flagellar signals appear and disappear periodically due to flagellar rotation. Yellow dots indicate the points at which each signal disappears. **d** Relationship between swimming speed and flagellar rotation rate (*N* = 44 cells). **e** Schematic illustration of TIRF microscopy. **f** Sequential images of flagellar filament rotation captured by TIRF microscopy. See also Movie S6. **g** Helix pitch of the flagellar filament bundle (*N* = 13 cells). **h** Model of anterior flagella-driven swimming in zoospore.

In *A*. *missouriensis*, predicted chemotaxis gene clusters are located at three loci (*che* clusters-1, −2, and −3)^17,23^. In a previous study, null mutants of each gene cluster (Δ*che*-1, Δ*che*-2, and Δ*che*-3 strains) were produced^17^. In this study, a null mutant strain of both *che*-1 and *che*-2 clusters (Δ*che*1–2) was generated by deleting *che* cluster-2 from the Δ*che*-1 chromosome, and a mutant strain lacking *fliC* and *che* clusters-1 and −2 (Δ*fliC*Δ*che*1–2) was generated by deleting *fliC* from the Δ*che*1–2 chromosome. We did not analyze *che* cluster-3 because this cluster does not appear to be involved in the chemotactic properties of zoospores, considering the predicted functions of gene products and transcriptional profiles during the life cycle^17^. We introduced the *fliC*^S260C^ gene into the Δ*fliC*Δ*che*1–2 strain using pTYM19-Apra to visualize flagellar filaments without chemotaxis signaling functions. Anterior localization was also observed in the Δ*fliC*Δ*che*1–2 strain harboring *fliC*^S260C^ (Movie S3), indicating that this configuration reflects the intrinsic propulsion architecture rather than a transient behavioral state. Together, these observations reveal that *A. missouriensis* zoospores swim using a front-positioned flagellar system, a configuration fundamentally distinct from the classical rear-driven bacterial swimming mode.

### High-speed swimming enabled by body-wrapping flagella

To resolve the propulsion mechanism directly, we visualized swimming zoospores using high-speed microscopy at 1,550 frames per second under laser illumination^24^. High-speed imaging revealed that a bundle of flagellar filaments wraps tightly around the anterior half of the cell body during swimming (Fig. 2b and Movie S4). Because the zoospores of the Δ*fliC*Δ*che*1–2 strain harboring *fliC*^S260C^ were used for this analysis, this configuration appears to correspond to the “run” mode during straight propulsion. Kymograph analysis showed wave-like propagation of the filaments away from the cell surface, with an apparent rotation frequency of ∼152 ± 33 Hz at 24°C (Fig. 2c). When the rotation rate and swimming speed were measured simultaneously in individual cells, the rotation frequency was nearly constant, whereas the swimming speed varied between 50 and 200 μm s^−1^ (Fig. 2d), consistent with propulsion driven by a coordinated filament bundle rather than stochastic filament activity. Imaging of immobilized, fluorescently labelled zoospores revealed approximately ten short filaments on one hemisphere (Fig. S2 and Movie S5), in agreement with values previously obtained by electron microscopy^15^, supporting the idea that these filaments rotate synchronously. Total internal reflection fluorescence (TIRF) microscopy of cells swimming near the coverslip further revealed that the filament bundle is oriented obliquely relative to the swimming axis (Fig. 2e-g and Movie S6). Because imaging was performed from the coverslip side without mirror inversion, these observations indicate that the flagellar bundle forms a right-handed (RH) helix around the cell body (Fig. 2h). These results demonstrate that zoospore propulsion is driven by a synchronously rotating, anterior, body-wrapping flagellar bundle forming an RH helix.

### Enhanced dispersal capacity of zoospores under low shear flow

Does ultrafast swimming confer a dispersal advantage under flow? To address this question, we examined zoospore migration in a laminar-flow chamber that generated a stable interface between a zoospore suspension and cell-free buffer (Fig. 3a). Under strong shear (5.9 mm s^−1^; 0.37 Pa), zoospores were passively advected by the flow. Strikingly, under weak flow (59 μm s^−1^; 3.7 × 10^−3^ Pa), zoospores rapidly crossed the laminar interface and invaded the cell-free region (Fig. 3b *left* and Movie S7), demonstrating active dispersal across streamlines. In contrast, *E. coli* cells showed negligible cross-interface migration under the same conditions (Fig. 3b *right*). Quantitative analysis showed that the apparent diffusion coefficient of zoospores under weak flow was ∼4,800 μm^2^ s^−1^, approximately 28-fold higher than that of *E. coli* (Fig. 3cd). These results indicate that zoospores retain substantial dispersal capability even in flowing environments. Hydrodynamically, rod-shaped bacteria tend to align with shear flow due to shape-induced torques, restricting their cross-stream transport^25^. The spherical geometry of zoospores likely minimizes such alignment, allowing more isotropic motion and facilitating efficient escape from streamlines (Fig. 3d *inset*). Together, these findings suggest that ultrafast swimming combined with spherical morphology enhances dispersal in heterogeneous flow environments.

**Figure 3:**
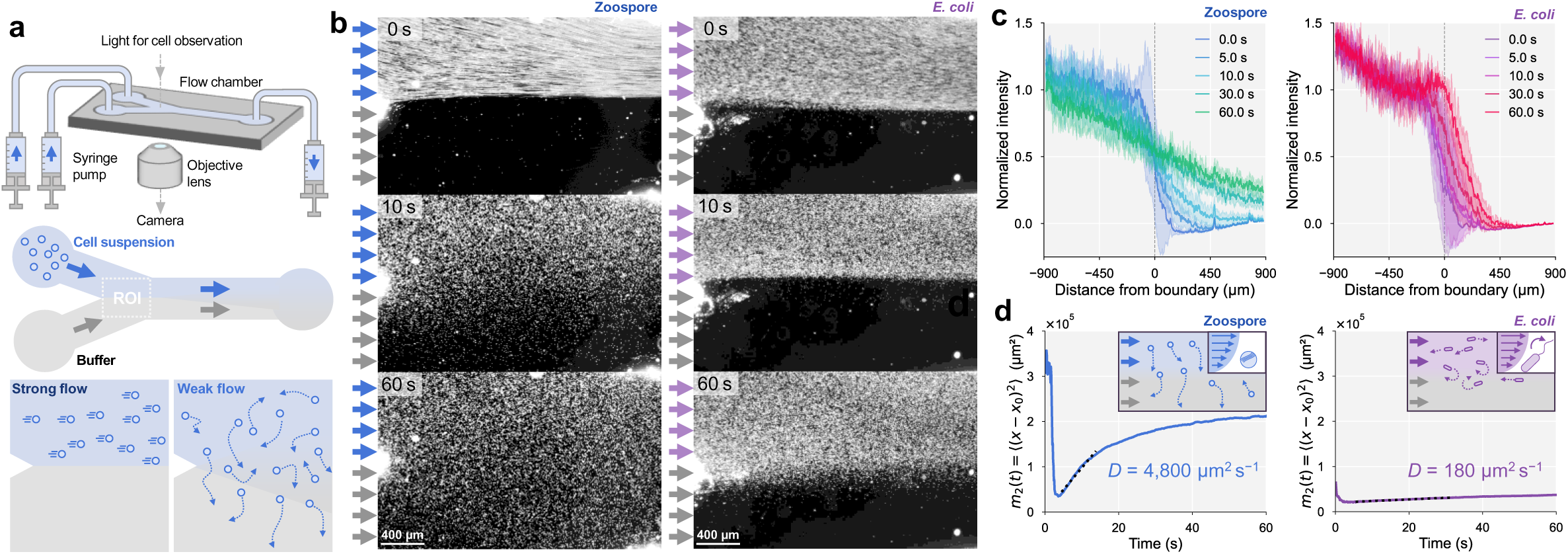
Enhanced dispersal of zoospores in a laminar flow chamber. **a** Experimental setup. *Top*: two solutions were introduced into the laminar-flow chamber using a syringe pump. *Middle*: bacterial suspension (upper inlet) and cell-free buffer (lower inlet) generate a stable laminar interface. *Bottom*: two-step flow protocol consisting of an initial high flow to remove cells followed by a low flow to observe dispersal. **b** Cell behaviors at the laminar interface captured under dark field microscopy. Representative images at 0 s, 10 s, and 60 s after reducing the flow speed for *A. missouriensis* zoospores and *E. coli* cells at 24°C. See also Movie S7. **c** Intensity profiles along a line perpendicular to the laminar boundary at different time points for zoospores (*left*) and *E. coli* (*right*). Intensities were normalized by setting the mean intensity on the cell-suspension side to 1 and that on the cell-free buffer side to 0. Independent replicate experiments: *N* = 5 for zoospores and *N* = 3 for *E. coli*. **d** Estimation of the apparent diffusion coefficient. The spatial spreading of the intensity profiles in panel **c** was quantified using the second moment of the distribution (0.1 s intervals), treating the intensity as a proxy for bacterial density. The apparent one-dimensional diffusion coefficient was estimated from the linear increase of the second moment over time. The initial transient regime was excluded, and the black dashed line represents the linear fit to the diffusive regime. *Inset*: schematic illustration of bacterial diffusion under weak flow.

### Chemotaxis and directional changes of zoospore swimming

Because it is known experientially that diluting zoospore suspensions reduces zoospore swimming speed, we tracked individual zoospores in 500-fold dilute suspensions to determine whether zoospore trajectories are actively regulated (Fig. 4a). Indeed, WT zoospores swam at a lower speed in dilute suspensions than in dense conditions (Fig. 1b and Fig. 4b), although the cause remains unclear. Interestingly, WT zoospores frequently followed curved paths punctuated by abrupt reorientation events, whereas Δ*che*1–2 zoospores swam along markedly straighter trajectories (Fig. 4a and Movie S8). In WT cells, directional changes exceeding 60° occurred within ∼0.6 s, while such rapid reorientations were rare in mutant cells (Fig. 4c). These results indicate that zoospores actively control their swimming direction through a *che*-dependent signaling pathway analogous to canonical bacterial chemotaxis systems. Remarkably, WT zoospores swam at approximately half the speed of Δ*che*1–2 zoospores (Fig. 4b). This result suggests that the swimming speed of zoospores is also controlled by a *che*-dependent signaling pathway. If this is the case, there may be a relationship between the regulatory mechanisms for reorientation and swimming speed. This interpretation suggests that directional control in zoospores is achieved not by altering propulsion architecture, but by modulating the activity of individual motors within the anterior flagellar arrays. Notably, chemotaxis-dependent reorientation of WT zoospores was most evident in dilute suspensions, whereas they appeared to favor persistent motion and dispersal over frequent steering in dense suspensions; however, the cause remains unclear. Together, these results indicate that the ultrafast anterior propulsion system can be dynamically regulated, enabling zoospores to balance rapid dispersal with directional control.

**Fig. 4:**
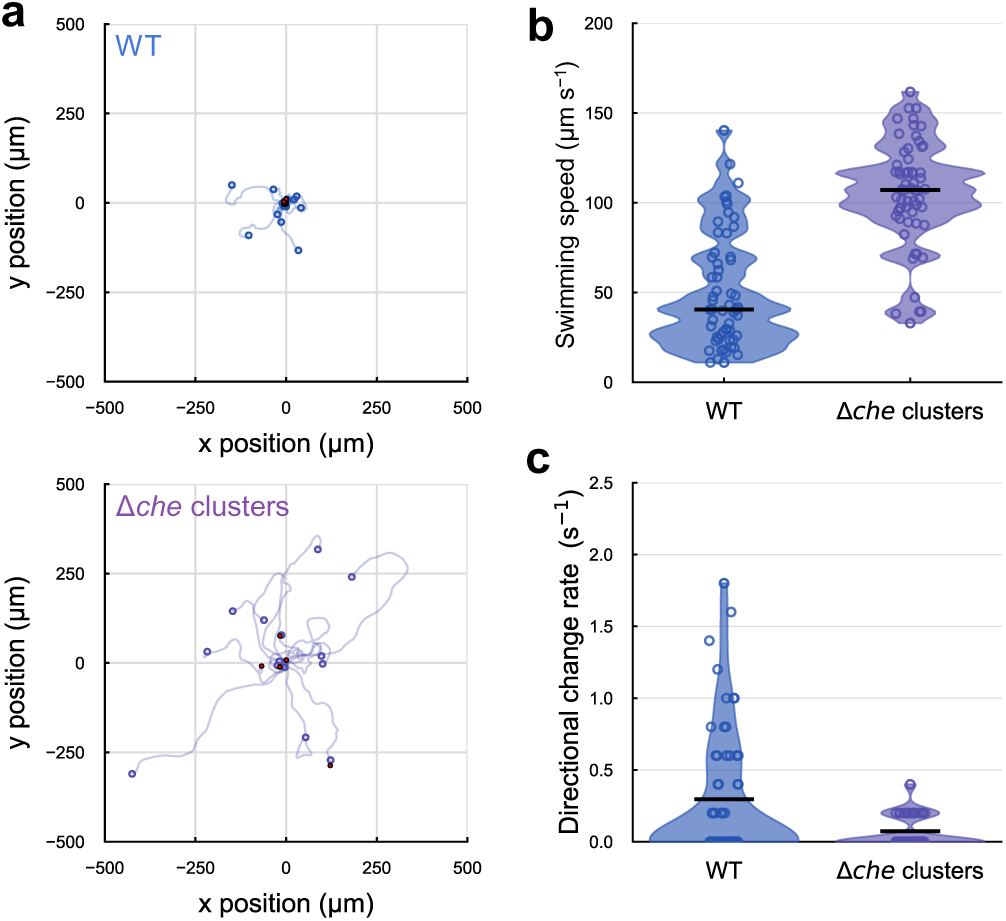
Directional change in zoospore swimming. **a** Single-cell trajectories of zoospores of the WT and Δ*che*1-2 strains recorded for 5 s in a dilution buffer at 24°C. The trajectories were smoothed using a Gaussian filter. Open circles indicate the endpoints of the trajectories. Red circles mark directional changes exceeding 60° within 0.6 s (*N* = 15 cells). **b** Swimming speed of cells moving faster than 10 µm s^−1^. Black bars indicate median values, and circles represent biological replicates (*N* = 60 cells). **c** Frequency of directional changes (>60° within 0.6 s) during swimming. The analysis included cells moving faster than 10 µm s^−1^. Black bars indicate mean values, and circles represent biological replicates (*N* = 60 cells).

## Discussion

Here we uncover an ultrafast swimming strategy in bacterial zoospores of *A. missouriensis*, establishing it as the fastest microorganism known when normalized by body length (Fig. 1). Their speeds exceed those reported for high-speed bacteria such as *Bdellovibrio bacteriovorus*^26^ and magnetotactic coccoid bacteria^8,9^, and approach those of fast-swimming hyperthermophilic archaea^10^. More importantly, our observations reveal not merely an increase in speed, but a distinct propulsion architecture based on an anterior, body-wrapping flagellar bundle.

Flagellar wrapping has previously been described as a transient behavioral state in polar-flagellated bacteria^27,28^, typically triggered by motor reversal and associated with directional switching^29,30^. In contrast, *A. missouriensis* zoospores exhibit wrapping even in mutants lacking chemotaxis gene clusters (Fig. 2), indicating that wrapping is a constitutive structural configuration rather than a transient response. Because the flagellar bundle forms an RH helix, propulsion is most consistent with counterclockwise (CCW) rotation of the bundle while maintaining the wrapped geometry. The short filaments adopt a tightly coiled morphology consistent with flagellar polymorphism models^31^ and with theoretical predictions of wrapping transmission^32^. This architecture presents an apparent paradox: in most bacteria, flagellar wrapping is thought to reduce propulsion efficiency in open fluid environments^29^. How, then, can zoospores achieve such extraordinary speeds? One possibility is that their shortened filaments and simplified basal rod; flagella of the phylum Actinobacteria, including *A. missouriensis*, lack the canonical rod proteins FlgF and FlgG^12^, representing evolutionary specializations that mechanically stabilize the wrapped bundle and enhance thrust transmission. Whereas wrapping in polar-flagellated bacteria has been linked primarily to motility in confined spaces^33^, our findings suggest that a wrapped, anterior bundle can instead function as a highly efficient propulsion module even in unconfined environments.

Our results suggest that ultrafast swimming in zoospores is achieved not by extreme motor speeds but by collective mechanical coordination of multiple flagellar filaments (Fig. 2). Whereas *Vibrio alginolyticus* attains high-speed swimming by rotating a single polar flagellum at up to 1,700 Hz^34^, *A. missouriensis* zoospores maintain comparatively moderate rotation rates at ∼150 Hz while deploying roughly a dozen filaments in a tightly coordinated bundle. Similar strategies are observed in other fast swimmers: hyperthermophilic archaea employ several dozen of filaments for propulsion^10,35^, and magnetotactic coccoid bacteria use compact multi-filament bundles enclosed within a sheath^8,9^. These systems point to a common physical principle in which propulsion force is amplified through filament number, spatial organization, and synchronous rotation rather than through increased motor speed alone. Such coordination may represent a form of collective synchronization at the microscale, optimizing torque transmission and energetic efficiency across multiple motors. From an evolutionary perspective, the anterior propulsion observed in zoospores may reflect a recurring solution for rapid dispersal (Fig. 3). The fastest known bacterium, *Candidatus* Ovobacter propellens, also positions its flagellar bundle at the front of the cell body^11^, suggesting that front-driven propulsion can evolve independently in phylogenetically distant lineages. In *A. missouriensis*, the motile phase lasts only an hour after spore release^13^, after which cells transition to a sessile growth stage. Under such constraints, maximizing swimming speed and dispersal range during this brief window is likely critical for colonizing favorable microhabitats.

Ecologically, ultrafast motility may enhance encounters with nutrient-rich substrates or symbiotic partners. In aquatic environments, motile bacteria exploit similar dynamics to interact with phytoplankton and organic particles^36^. Analogously, zoospores in water temporarily held in the soil may be guided toward fungal hyphae, plant debris, or other nutrient sources through chemotaxis^37^. Notably, *A. misouriensis* has a large number of chemotaxis receptors; 21 methyl-accepting chemotaxis proteins (MCPs) are encoded in its genome. Although the mechanisms that regulate the motility of zoospores to exhibit chemotaxis are unknown, we observed clear differences between wild-type and Δ*che*1–2 zoospores in swimming speed and the frequency of abrupt reorientation events when the zoospore suspension was 500-fold diluted, providing clues toward elucidating these mechanisms. Our microfluidic experiments further showed that spherical zoospore cells disperse efficiently even under weak flow (Fig. 3), suggesting their morphology and propulsion mode together facilitate escape from streamlines and promote colonization in heterogeneous environments^38^. More broadly, the propulsion strategy revealed here, an anterior, body-wrapping bundle of synchronously rotating filaments, defines a previously unrecognized design principle for microscale locomotion. By achieving high speeds without extreme motor frequencies, this architecture may provide inspiration for bio-inspired micromachines capable of rapid navigation.

## Methods

### Strain and culture conditions

*A. missouriensis* 431^T^ (NBRC 102363^T^) was obtained from the National Institute of Technology and Evaluation (NITE), Chiba, Japan. *A*. *missouriensis* was grown on YBNM (yeast extract-meat extract-NZ amine-maltose) or HAT (humic acid-trace element) agar at 30°C for a solid culture and in PYM (peptone-yeast extract-MgSO_4_) broth at 30°C for a liquid culture, as described previously^19^. *E. coli* W3110 was used in the experiment using laminar flow chamber.

### Construction of deletion mutants and complemented strains

For construction of the Δ*che1*-2 and Δ*fliC*Δ*che*1-2 strains, the regions upstream and downstream of the *che* cluster-2 and *fliC* (approximately 2 kbp each) were amplified by PCR. The amplified DNA fragments were digested with restriction enzymes (Table S1) and cloned into pUC19 digested with the same restriction enzymes. The generated plasmids were sequenced to confirm that no PCR-derived error was introduced. The DNA fragments were digested with restriction enzymes and cloned together into pK19mobsacB, in which the kanamycin resistance gene had been replaced with the apramycin resistance gene *aac(3)IV*^20,39^. The plasmids for the deletion of *che* cluster-2 and *fliC* were introduced into the *A. missouriensis* Δ*che*-1 and Δ*che*1-2 strains, respectively, by conjugation with *E*. *coli* ET12567 (pUZ8002). Apramycin-resistant colonies resulting from a single-crossover recombination were isolated. One of them was cultivated in PYM liquid medium at 30°C for 36–48 h, and the mycelia were spread on the Czapek-Dox agar medium containing extra sucrose (final concentration of 5%). After incubation at 30°C for 5–7 days, sucrose-resistant colonies were inoculated on YBNM agar with or without apramycin to confirm their sensitivity to apramycin. Apramycin-sensitive and sucrose-resistant colonies resulting from a second single-crossover recombination were isolated as candidates for the gene deletion mutants. The disruption of *che* cluster-2 and *fliC* in the Δ*che*1-2 and Δ*fliC*Δ*che*1-2 strains, respectively, was confirmed by PCR using genomic DNA prepared from the candidate strains as a template. A 1.6-kbp DNA fragment containing the promoter and cysteine-substituted (S260C) coding sequences of *fliC* was amplified by overlap extension PCR. The amplified fragment was digested with restriction enzymes (Table S1) and cloned into pTYM19-Apra^20,21^. The generated plasmid was sequenced to confirm that no PCR-derived error was introduced and introduced into the Δ*fliC* and Δ*fliC*Δ*che*1-2 strains by conjugation. The Δ*fliC* strain was obtained previously^16^. Apramycin-resistant colonies were then obtained as the *fliC*^S260C^-introduced mutant strains.

### Measurements of swimming speed

Zoospores were released from the sporangia by pouring 10 ml of 25 mM NH_4_HCO_3_ onto one HAT plate and incubating the plate at 30°C for 1 h. The liquid was collected from the plate using a pipette as a zoospore suspension. The sample chamber was constructed by sandwiching two strips of double-sided tape between a glass slide and coverslip to form a narrow channel. The chamber was pre-coated with BSA buffer containing 2% (wt/vol) BSA in 25 mM NH_4_HCO_3_. The above-mentioned zoospore suspension was introduced into the chamber, and both ends were sealed with nail polish to keep the sample from drying. For dilution-based observations for chemotaxis, the suspension was diluted 500-fold in 25 mM NH₄HCO₃ prior to imaging.

### Laminar flow chamber

The flow chamber was assembled by taping a coverslip with a drilled glass slide, as described previously^40^. Inlet and outlet holes (1.0 mm diameter) were bored with a high-speed drill with a diamond-tipped bit (No. 13853; NAKANISHI). A Y-shaped channel was cut into 0.1 mm-thick double-sided tape (7082, Teraoka) with a bifurcation angle of 50 degree. Each branched arm measured 15 mm in length and 2.0 mm in width, and the main channel measured 31 mm in length and 2.0 mm in width. Inlet and outlet ports (N-333; IDEX Health & Science) were attached with elastic adhesive (Super X No. 8008 clear, CEMEDINE). A syringe pump (Legato 200, Kd Scientific) was connected to the flow chamber by a connecter and tube (F-333NX and 1512 L, IDEX Health & Science). The flow chamber was pre-coated with BSA buffer. A zoospore suspension and an *E. coli* cell suspension were simultaneously introduced through the inlet channels. A strong flow (1 µL/s) was applied for 20 s, followed by a weak flow (10 nL/s); both the flow rates and their transition were programmed. Image analysis was conducted after the flow stabilized following the change in flow rate and after confirming that no bacteria had diffused across the boundary region.

### Quantification of bacterial spreading and estimation of the apparent diffusion coefficient

Bacterial spreading in the microchannel was quantified from the intensity profile along a line perpendicular to the laminar boundary, using the Plot Profile function in ImageJ (line width = 1200 pixels, 176 µm). The intensity signal was used as a proxy for bacterial density. To remove the background signal, the initial intensity profile was subtracted:

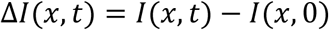

Negative values resulting from noise were set to zero. The analysis was restricted to the cell-free buffer region beyond the boundary position *x*_0_, where no bacteria were initially present. The spatial spreading of the bacterial population was quantified using the second moment of the intensity distribution^41^:

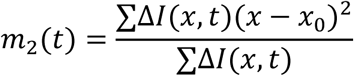

where *x* − *x*_0_represents the distance from the boundary. For one-dimensional diffusive spreading, the second moment increases linearly with time. The apparent diffusion coefficient, *D*_app_, was therefore estimated from

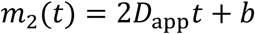

where *b* is a constant offset. Linear regression of *m*_2_(*t*) yielded the slope *a*, from which the apparent diffusion coefficient was calculated as

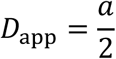

To avoid early-time noise and late-time saturation, fitting was restricted to time points at which the total signal mass *M*(*t*) was between 10% and 50% of its maximum value.

### Preparation of fluorescently labeled zoospore

Flagellar filaments of cysteine-substitution mutants were labeled using DyLight 488 maleimide-conjugated fluorophores (Thermo-Fisher). Zoospores were released from the sporangia by pouring 10 mL of 25 mM NH_4_HCO_3_ onto one HAT plate at 30°C for 1 h. DyLight dye (10 μg/μL in DMSO) was added directly to the plate at a volume of 10 μL at the onset of sporangium rupture. The suspension of labeled zoospore was centrifuged at 5,000 × g for 2 min at 25 °C, washed twice, and resuspended in washing buffer. The washing buffer was prepared by overlaying 10 mL of fresh 25 mM NH_4_HCO_3_ onto a cell-free HAT plate and incubating it for 1 h. After washing, the zoospore suspension was used for the observation within 1 h.

### Optical microscopy

Swimming zoospore cells were visualized under a dark-filed microscope (IX73, Olympus) equipped with an objective lens (UCPLFLN20×, NA 0.70, and UPLFLN4×PH-2, NA 0.13, Olympus), a CMOS camera (DMK 33UX174, Imaging Source), and an optical table (HAX-0806, JVI). The microscope stage was heated with a thermo-plate (TP-110R-100; Tokai Hit). Projections of the images were captured as greyscale images with the camera under 10-ms, 5-ms, and 4-ms intervals and converted into a sequential TIF file without any compression. All data were analyzed by ImageJ 1.53k (rsb.info.nih.gov/ij/) and its plugins, TrackMate^42^, and particle tracker^43^. For direct visualization of fluorescently labeled zoospore, the sample was examined under an inverted microscope equipped with an objective lens (UCPLFLN20×, NA 0.70, or UPLXAPO100×OPH, NA 1.45, Olympus), a dichroic mirror (Di02-R488, Semrock), dual-view imaging system with optical filters (FF560-FDi01, FF03-535/50 and BLP01-568R, Semrock), a CMOS camera (Zyla 4.2, Andor), and an optical table (ASD-1510T, JVI). A laser beam (488-nm wavelength, OBIS488, Coherent) was introduced into the inverted microscope through an objective lens side. Projections of the images were captured with the camera under 5-ms or 0.625-ms intervals for free-swimming cells under epi-illumination and TIRF and converted into a sequential TIF file without any compression.

### Electron microscopy

The *A*. *missouriensis* WT strain was cultivated on HAT agar at 30°C for 7 days to induce sporangium formation. The release of spores was induced by pouring the 25 mM NH_4_HCO_3_ solution onto the sporangium-forming HAT agar plate (10 ml per plate) and incubating the plate at room temperature for 1 h. The zoospore suspension was retrieved from the agar surface, and the zoospores bound to the carbon-coated grids (Nisshin-EM, Tokyo, Japan) were stained with 1% (wt/vol) phosphotungstic acid solution (pH 7.0) and air-dried. Samples were observed under a JEM-1010 or JEM-1400Plus transmission electron microscope (JEOL, Tokyo, Japan) at 80 kV.

### Data availability

All data are available in the main text and the Supplementary Information/Source data file with this paper.

## Acknowledgments

We thank Bo Jiang for her technical support in generating recombinant *A. missouriensis* strains. This study was supported partly by KAKENHI grants from the Japan Society for the Promotion of Science (JSPS) 22H05066 to DN, 17K07711 to TT, 18H02122 and 24H00500 to YO and FOREST Program from Japan Science and Technology Agency (JST) JPMJFR2411 to DN. The funders had no role in study design, data collection and analysis, decision to publish, or preparation of the manuscript.

## Author contributions

Conceptualization: TT, DN and YO, Methodology: NAU, TI, YS, MSJ, TT and DN, Formal analysis and investigation: NAU and DN, Funding acquisition: TT, DN and YO, Writing – original draft: NAU, TT, DN and YO, Writing – review & editing: TT and DN.

## Supplementary Information

### Supplementary Figures

**Fig. S1:**
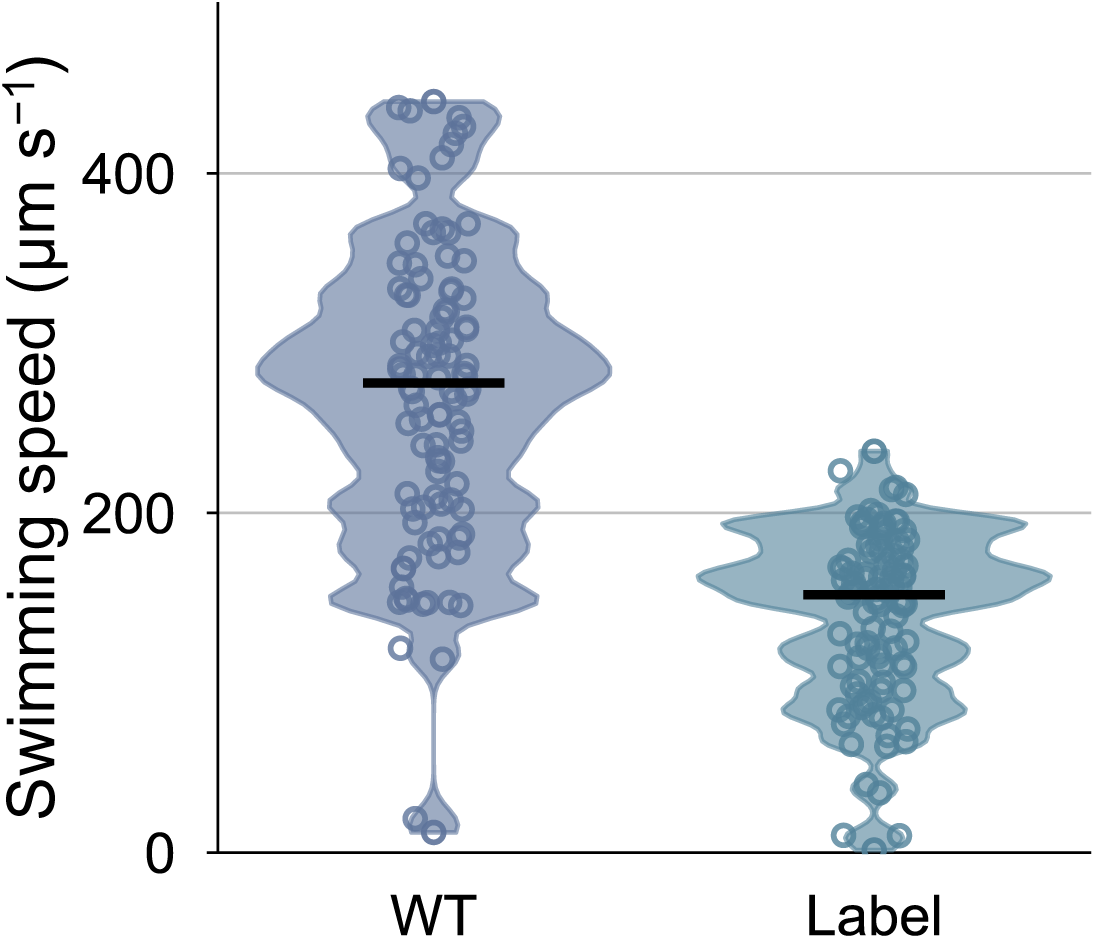
Swimming speed of flagellar filament-labeled cells. Swimming speed of *fliC*^S260C^ mutant zoospores at 24°C. Black bars indicate the median values, and circles represent biological replicates (*N* = 100 cells).

**Fig. S2:**
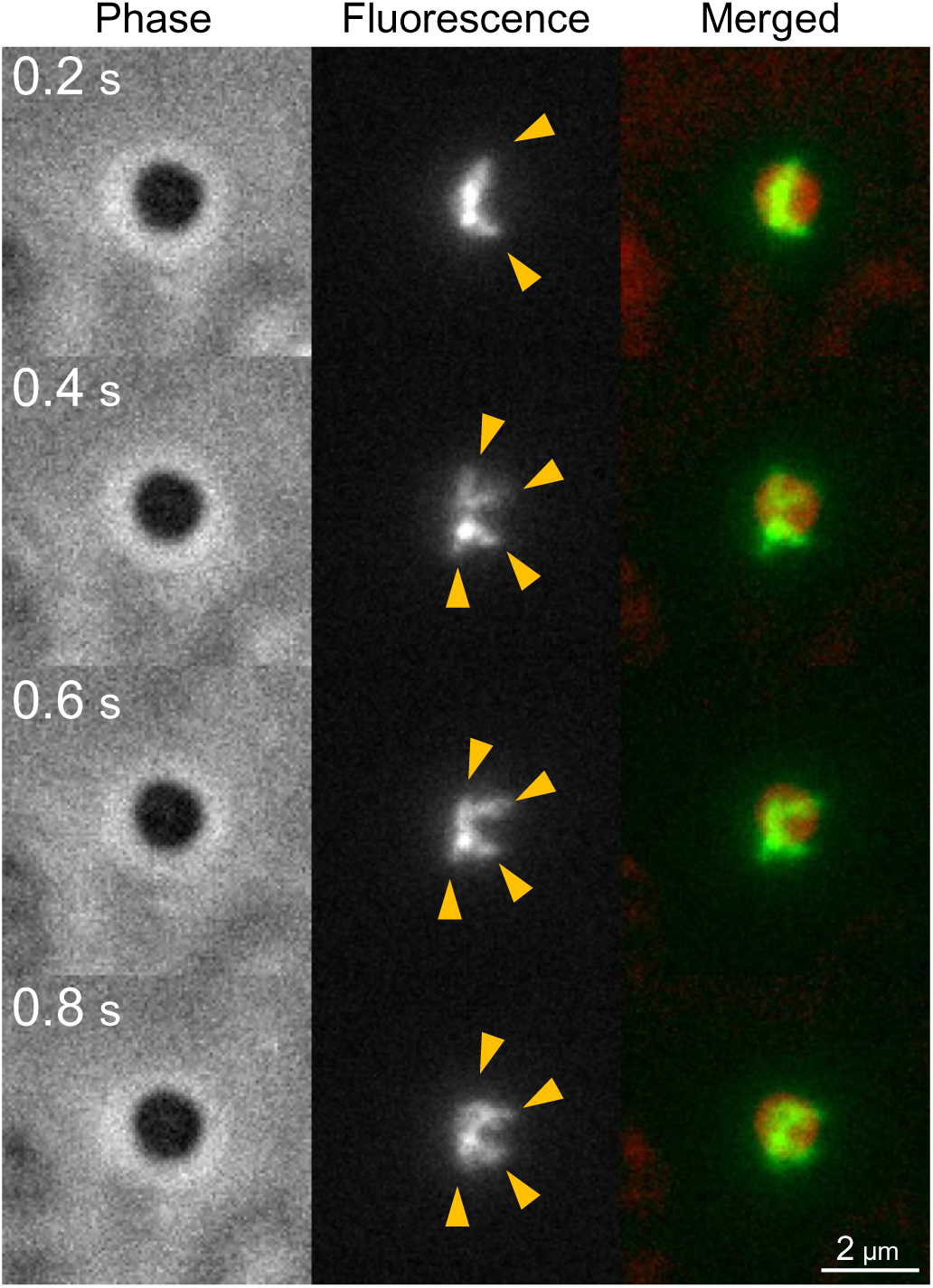
Visualization of flagellar filaments in an immobilized zoospore. A time-lapse series of an immobilized cell. Phase-contrast image (*left*), fluorescence image of labeled flagellar filaments (*middle*), and merged image (*right*). Orange arrowheads indicate flagellar filaments. See also Movie S5.

### Supplementary Table

**Table S1.**
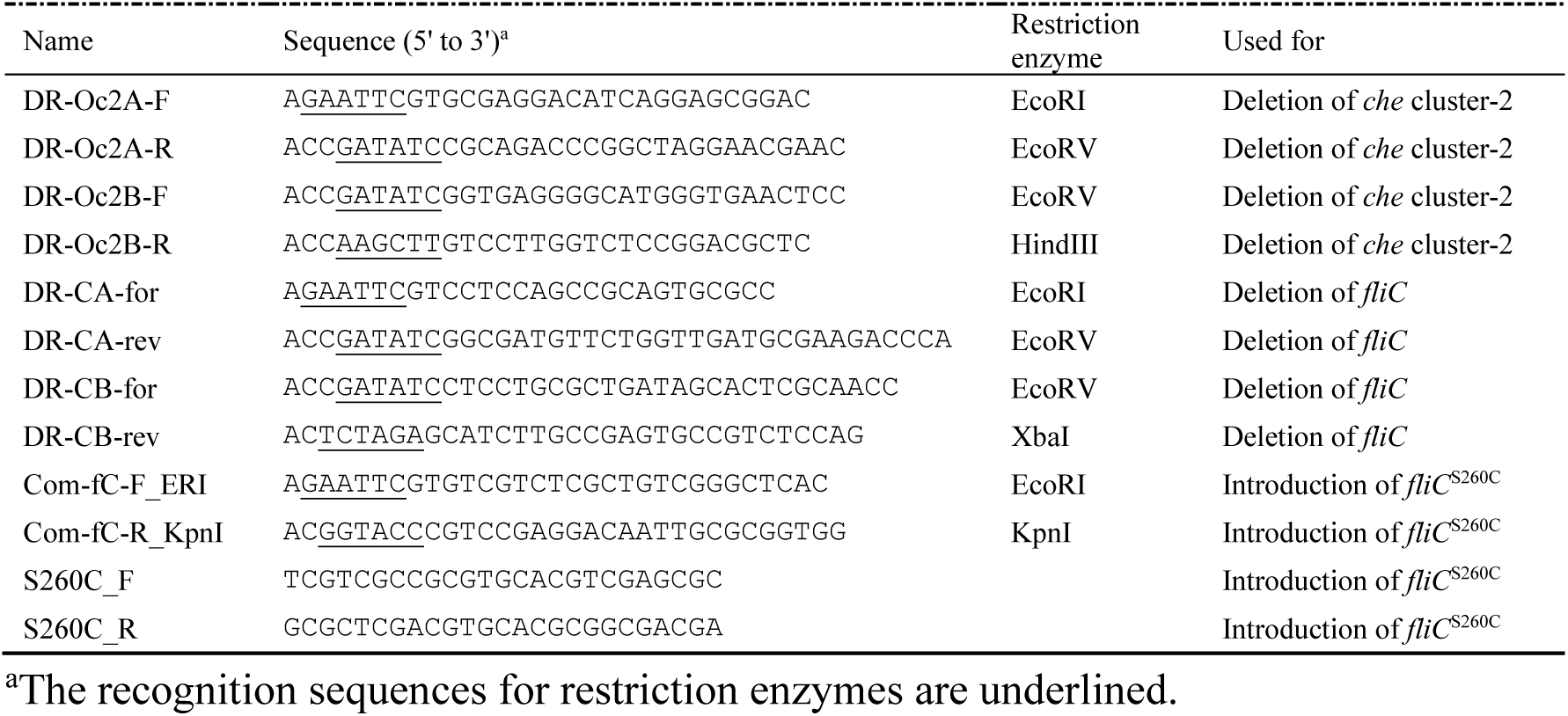
Oligonucleotides used in this study.

## Legend for Supplementary Movies

**Movie S1. Release of zoospores from a sporangium.**

Dark-field microscopy showing the release of zoospores from a sporangium. In the region marked by the blue box, a zoospore is released at ∼8 s and immediately begins rapid swimming. In the second half of the movie, this region is magnified to show the zoospore of interest indicated by an arrowhead, and the release event is shown in slow motion.

**Movie S2. Ultrafast swimming of zoospores.**

Zoospore swimming observed under dark-field microscopy at 30°C, the optimal temperature for mycelial growth of this bacterium. The movie shows the original recording, the tracked trajectories, and a slow-motion version of the tracked trajectories. Swimming paths are color-coded according to instantaneous speed, following the scale shown in the lower-left corner.

**Movie S3. Zoospores swim with anteriorly positioned flagella.**

Flagellar filaments were fluorescently labeled and observed during swimming at 10 ms intervals. Zoospores were visualized by dark-field microscopy (red channel), while flagellar filaments were detected by fluorescence microscopy (green channel). The two channels were simultaneously projected onto a single camera sensor using a dual view system. Zoospores with *fliC* cysteine substitution mutant in the background of WT (left) and Δ*che*1–2 (right) are shown. The movie shows real-time playback followed by slow motion.

**Movie S4**. **High-speed imaging of flagellar rotation in swimming zoospores**

Flagellar filaments of the *fliC* cysteine substitution mutant in Δ*che*1–2 background were fluorescently labeled and imaged at 0.625 ms intervals using high-speed fluorescence microscopy. Area, 7.8 μm × 5.9 μm.

**Movie S5. Flagellar dynamics in an immobilized zoospore.**

Simultaneous observation of the cell body and flagellar filaments using a dual view system. Zoospores and flagellar filaments were imaged by phase-contrast microscopy and fluorescence microscopy, respectively. Left: phase contrast. Center: fluorescence. Right: merged image. Area, 6.5 μm × 19.5 μm.

**Movie S6. High-speed TIRF imaging of swimming zoospores.**

TIRF microscopy of a swimming zoospore with fluorescently labeled flagellar filaments (*fliC* cysteine substitution strain in the Δ*che*1–2 background). Images were acquired at 0.625 ms intervals. Cell bodies and flagella were visualized by phase contrast microscopy and fluorescence microscopy, respectively. Left: phase contrast. Center: fluorescence. Right: merged image. Area, 4.9 μm × 19.5 μm.

**Movie S7**. **Rapid dispersal of zoospores in a laminar-flow environment**

Laminar flow was generated in a microfluidic chamber by introducing two solutions using syringe pumps: a cell suspension in the upper layer and a cell-free buffer in the lower layer. When the flow velocity was reduced to 59 µm/s, zoospores crossed the interface and dispersed into the lower layer. Left: zoospores. Right: *E. coli*. Dispersals were observed under dark-field microscopy for 1 min.

**Movie S8**. **Chemotaxis-dependent reorientation of zoospore swimming**

Swimming trajectories of zoospores in dilute suspension observed under dark-field microscopy. Left: WT. Right: Δ*che*1–2 mutant. The movie shows the original recording and the tracked trajectories. WT cells frequently exhibit abrupt directional changes, whereas the mutant shows predominantly straight swimming trajectories.

